# Characterization of β-Hydroxybutyrate as a Cell Autonomous Fuel for Active Excitatory and Inhibitory Neurons

**DOI:** 10.1101/2024.06.08.598077

**Authors:** Kirsten Bredvik, Charmaine Liu, Timothy A. Ryan

## Abstract

The ketogenic diet is an effective treatment for drug-resistant epilepsy, but the therapeutic mechanisms are poorly understood. Although ketones are able to fuel the brain, it is not known whether ketones are directly metabolized by neurons on a time scale sufficiently rapid to fuel the bioenergetic demands of sustained synaptic transmission. Here, we show that nerve terminals can use the ketone β-hydroxybutyrate in a cell- autonomous fashion to support neurotransmission in both excitatory and inhibitory nerve terminals and that this flexibility relies on Ca^2+^ dependent upregulation of mitochondrial metabolism. Using a genetically encoded ATP sensor, we show that inhibitory axons fueled by ketones sustain much higher ATP levels under steady state conditions than excitatory axons, but that the kinetics of ATP production following activity are slower when using ketones as fuel compared to lactate/pyruvate for both excitatory and inhibitory neurons.

**Significance Statement:** The ketogenic diet is a standard treatment for drug resistant epilepsy, but the mechanism of treatment efficacy is largely unknown. Changes to excitatory and inhibitory balance is one hypothesized mechanism. Here, we determine that ATP levels are differentially higher in inhibitory neurons compared to excitatory neurons, suggesting that greater mitochondrial ATP production in inhibitory neurons could be one mechanism mediating therapeutic benefit. Further, our studies of ketone metabolism by synaptic mitochondria should inform management of side effects and risks associated with ketogenic diet treatments. These results provide novel insights that clarify the role of ketones at the cellular level in ketogenic diet treatment for intractable epilepsy and inform the use of ketogenic diets for neurologic and psychiatric conditions more broadly.

## Introduction

The body’s homeostatic mechanisms typically conserve glucose for use by the brain, but during glucose starvation the brain may use ketones as a mitochondrial fuel (Zhang et al., 2015). Active neurons can adapt to non-glycolytic fuel sources in real time to match continuously fluctuating energetic demands (Rangaraju et al., 2014; Pathak et al., 2015; Ashrafi et al., 2020; Zampese et al., 2022), but whether neurons independently metabolize ketones is not known. Besides the standard mitochondrial fuel of pyruvate, there is evidence that alternative carbon sources such as amino acids and glycerol can be metabolized by mitochondria to support neuronal function (Divakaruni et al., 2017; Dhoundiyal et al., 2022). Ketone bodies are a notable example of an alternative mitochondrial fuel, as the ketogenic diet is in use as a therapy for medication-refractory cases of epilepsy (El-Rashidy et al., 2023) and is effective in about 40% of these cases (Ye et al., 2015). There are multiple proposed mechanisms for the therapeutic effect of the ketogenic diet including epigenetic modifications, neuroprotective signaling, antioxidant activity, and metabolic shifts leading to shifts in the ratio of excitation to inhibition (Murugan and Boison, 2020; Ma et al., 2007; Juge et al., 2010; Qiao et al., 2024), each with varying amounts of supporting evidence. The efficacy of the ketogenic diet is relatively high for inherited metabolic disorders with impaired glycolysis (Klepper et al., 2005; Sofou et al., 2017), supporting the notion that an underlying metabolic component contributes to the therapeutic impact. Recent broader applications of the ketogenic diet to modulate neuronal activity in neurodegeneration (Ramezani et al., 2023) and psychiatric conditions (Sethi et al., 2024) raise the possibility that metabolism could modulate synaptic activity and/or circuit dynamics even in non-epileptic individuals.

It remains to be demonstrated conclusively that neurons express and effectively employ the necessary enzymes to break down ketones and use them to produce ATP. For the metabolism of ketones to plausibly contribute to the antiepileptic effect of the ketogenic diet, it is important to determine to what extent excitatory and inhibitory neurons can perform ketone metabolism and whether this metabolism can occur at a sufficient rate to support active neuron function. Understanding ketone metabolism and its regulation and limitations in both excitatory and inhibitory neurons will provide key insights into the fundamental functions of mitochondria in neurons and the influences of metabolism on neural circuit function.

Differential expression of genes and proteins involved in mitochondrial metabolism suggests that inhibitory neurons in aggregate may be better equipped to make use of mitochondrial fuel sources than excitatory neurons (Nie et al., 1995; Gulyás et al., 2006; Fecher et al., 2019; Wynne et al., 2021). Inherently faster-spiking inhibitory neurons have unique ATP requirements (Inan et al., 2016; Kann et al., 2011) that may require different regulatory mechanisms than excitatory neurons to supply sufficient ATP for synaptic transmission. While neurons upregulate mitochondrial metabolism through Ca^2+^ signaling during activity (Llorente-Folch et al., 2015), this upregulation could occur through activation of PDH or dehydrogenases in the tricarboxylic acid cycle (Denton, 2009). Evidence suggests that pyruvate dehydrogenase (PDH) is a key regulated step in the metabolism of highly active neurons (Yan et al., 2024), but it is not known whether an alternative mitochondrial fuel that bypasses PDH could robustly support the large metabolic burden (Rangaraju et al., 2014) of neuronal firing. As a potential neuronal fuel that bypasses PDH, ketones present a unique opportunity to study differential metabolic regulation by Ca^2+^ in the mitochondria of excitatory and inhibitory neurons.

We here employ cell type specific fluorescent reporters in hippocampal excitatory and inhibitory neurons in parallel to compare their ability to metabolize the ketone body β-hydroxybutyrate (BHB). We find that neurons are able to use BHB to fuel synaptic transmission in a cell autonomous manner, that Ca^2+^-dependent upregulation of mitochondrial metabolism is necessary to utilize either BHB or pyruvate as fuel for synaptic transmission.

## Materials and Methods

### Animals

Wild type Sprague-Dawley rats at age 1 to 3 days old were used for all experiments (Charles River code 400, RRID: RGD_734476). Mixed gender rat were sacrificed and dissected in accordance with protocols approved by Weill Cornell Medicine IACUC.

### Culture and Transfection

Neurons were dissected from the CA1 to CA3 regions of the hippocampus, or from the hippocampus and cortex combined. Neurons were cultured and plated on glass coverslips coated with poly-L-ornithine as described previously (Farrell et al., 2023). Calcium phosphate-mediated gene transfer was performed using a one hour incubation in order to transfect 6- to 7-day-old cultures as described previously (Sankaranarayanan et al., 2000). Neurons were incubated at 37° C in a 95% air/5% CO_2_ humidified incubator, and then imaged 14-21 days after plating. Neuronal culture media consisted of MEM (Thermo Fisher #51200038), 0.25 g/L insulin, 0.3 g/L GlutaMAX, 5% fetal bovine serum (Atlanta Biological S11510), 2% N-21 (Chen et al., 2008), and 4 µM cytosine β-d-arabinofuranoside.

### Western blots

Cortical neurons cultured from 1- to 3-day-old rats were plated in 6-well plates that had been coated with poly-L-ornithine. After 2-3 days, cytosine β-d-arabinofuranoside was added to the media using the same media formulations as for hippocampal cultures (Farrell et al., 2023). Approximately 500,000 GC of lentivirus was added to a well containing about 2.5 million cells including a mixture of neurons and glia. After 10-11 days of incubation with lentivirus cultures were harvested and prepared for western blot using RIPA buffer (Thermo Fisher, 89900). Samples were heated for 10 minutes at 95° C with 50 mM DL-dithiothreitol (Sigma Aldrich, 43819) and Laemmli buffer at 1X (Bio-Rad, 161-0747) and loaded into 4-20% precast gels (Bio-Rad, 4561096 and 4561094). After running for 45 minutes at 200 V in Tris/Glycine/SDS buffer (Bio-Rad, 1610732), gels were transferred to nitrocellulose membrane (Bio-Rad, #1620115) at 100 V for 30 minutes at 4° C in Tris/Glycine buffer (Bior-Rad, 1610734) using 20% methanol. Total protein on the gel was quantified with REVERT™ total protein stain (LI-COR, #926-11011), using manufacturer instructions. Membranes were blocked for one hour with Odyssey Blocking Buffer (TBS) (LI-COR, #927-50000) and incubated overnight at 4° C with primary antibody in Odyssey Blocking Buffer. For SCOT knockdown validation, OXCT1 Rabbit polyclonal antibody (Proteintech 12175-1-AP) was used at 1:2000. For BDH1 knockdown validation, rabbit BDH1 polyclonal antibody (Invitrogen PA5-56598) was used at 1:800. Secondary antibody (IRDye^®^ 800CW Goat Anti-Rabbit, LI-COR #926-32211) was incubated for one hour at room temperature, diluted 1:20,000 in Odyssey Blocking Buffer after washing with TBS + 0.1% Tween-20. LI-COR Odyssey^®^ Fc Imaging System and Image Studio Software was used to develop and quantify blots.

### Analysis

Western blots were analyzed with the “draw rectangles” function in LI-COR Image Studio Lite software. The signal of each lane was normalized to the total protein signal (REVERT™ total protein stain; Extended Data Figures 1-1, 1-3, 1-5, 1-7, S1-1, S1-3). For graphs, each replicate was additionally normalized to the control average.

Live images were analyzed with the Time Series Analyzer plugin in ImageJ, using 20-149 ROIs of ∼2 μm diameter to quantify the fluorescence of the synaptic boutons over time. To quantify change in fluorescence at a specific time point, the fluorescence of three frames were averaged and compared to a pre-stimulation baseline of between 10 and 25 frames. Maximal fluorescence change was quantified at the end of stimulation. iATPSnFR2.0-HALO measurements were taken using alternating frames of 488 and 637 laser illumination, where the cell averages of regions from individual frames imaged with a 488 laser were normalized to the paired cell average of ten frames imaged with a 637 laser prior to stimulation. For physin-pHluorin assay, responses where < 4% of vesicle pool was released within 5 seconds of stimulation were excluded from analysis. Whenever physin-pHluorin fluorescence traces were normalized to maximum in traces, raw change in fluorescence as proportion of maximal fluorescence in NH_4_Cl was also reported in Supplementary Table 1 (except for traces in Figure 1A and 1B, reported in source data only due to spatial constraints).

**Figure 1.**
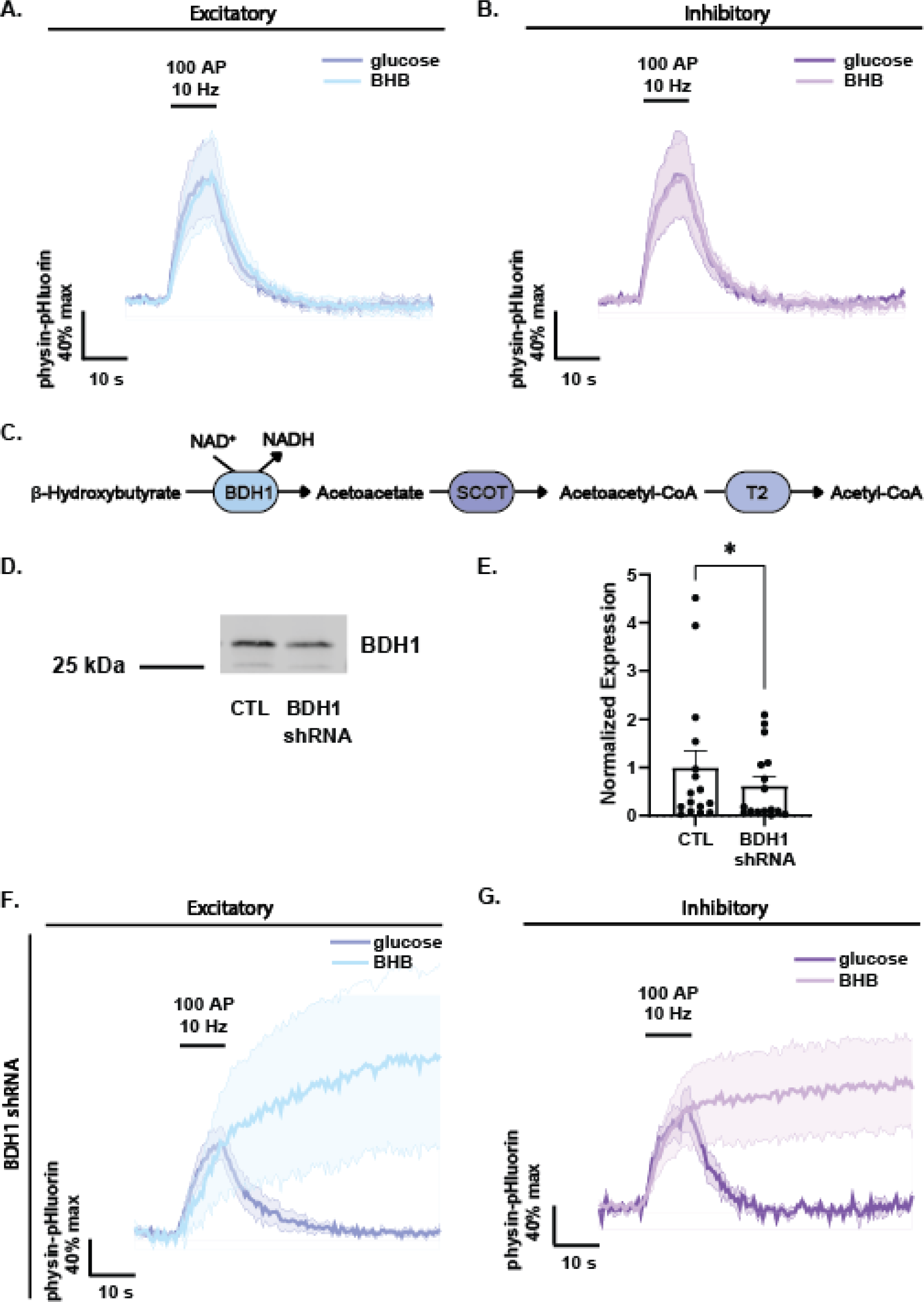
Neurons cell autonomously process the ketone β-hydroxybutyrate as fuel. **A)** Synaptic vesicle recycling following 100 action potential (AP), 10 Hz stimulus in excitatory neurons expressing CaMKII synaptophysin-pHluorin (physin-pHluorin), while fueled by 5 mM glucose or 5 mM β-hydroxybutyrate (BHB) (n=9, error bands SEM). **B)** Synaptic vesicle cycling in inhibitory neurons expressing hDlxI56i physin-pHluorin and fueled by glucose or BHB during 100 AP, 10 Hz stimulus (n=4, error bands SEM). **C)** Schematic of ketone catabolism in neurons by β-hydroxybutyrate dehydrogenase 1 (BDH1), succinyl-CoA:3-ketoacid CoA transferase (SCOT), and thiolase II (T2). **D)** Representative Western blot against BDH1 in WT cortical culture or culture treated with BDH1 shRNA virus for 10 days (full blots in Extended Data Figure 1-1, -2, -3, -4, -5, -6, -7, -8). **E)** Quantification of BDH1 knockdown efficiency in Western blot, normalized to average signal of uninfected controls (N=4 cultures, n=4 technical replicates, mean=0.6238. Error bars SEM, *p=0.011, Wilcoxon matched-pairs signed rank test). **F)** Synaptic vesicle recycling following 100 AP, 10 Hz stimulus in excitatory neurons expressing physin-pHluorin and shRNA against BDH1, while fueled by 5 mM glucose or BHB (n=4, error bands SEM). **G)** Synaptic vesicle cycling in inhibitory neurons expressing physin-pHluorin with and without shRNA against BDH1, while fueled by glucose or BHB during 100 AP, 10 Hz stimulus (n=4, error bands SEM).

### Imaging

For iATPSnFR2.0 experiments, cells were incubated with 50 nM Janelia Fluor® 635 HaloTag® ligand (JF-635) dye for 30 minutes, then washed with Tyrode’s buffer prior to imaging. Live imaging experiments were performed on a custom-built laser illuminated epifluorescence microscope using an Andor iXon+ camera (model DU-897U-CS0-#BVF). Coherent OBIS 488 nm and 637 nm lasers were used to illuminate samples and were controlled by an Arduino program. Images were acquired through a 40X 1.3 NA Fluar Zeiss objective. Coverslips were mounted in a laminar flow perfusion chamber and perfused with Tyrode’s buffer containing (in mM) 119 NaCl, 2.5 KCl, 2 CaCl_2_, 2 MgCl_2_, 50 HEPES (pH 7.4), 5 glucose or 5 sodium D-/L-β-hydroxybutyrate (Sigma) or 1.25 sodium lactate (Sigma) and 1.25 sodium pyruvate (Sigma), supplemented with 10 μM 6-cyano-7-nitroquinoxalibe-2, 3-dione (CNQX), and 50 μM D, L-2-amino-5-phosphonovaleric acid (APV) (both from Sigma-Aldrich) to prevent excitatory post-synaptic responses. Koningic acid (heptilidic acid; Cayman Chemical) was included at 10 μM in specified experiments. In some experiments, data from conditions of 5 mM β-hydroxybutyrate and 5 mM glucose + koningic acid + 5 mM β-hydroxybutyrate were pooled, as these conditions were assumed to be metabolically similar. When NH_4_Cl solution was used for pHluorin-containing vesicle alkalization, it contained (in mM) 50 NH_4_Cl, 69 mM NaCl, 2.5 KCl, 2 CaCl_2_, 2 MgCl_2_, 55 HEPES (pH 7.4).

Action potentials were evoked in neurons by delivering 1 ms pulses between platinum–iridium electrodes, resulting in an electric field potential of ∼10 V/cm. Stimulation timing was controlled using custom-built Arduino and python programs. Temperature was maintained at 37°C with a custom-built objective heating jacket. For live imaging of knockdown neurons, co-transfection of shRNA and pHluorin plasmids was confirmed visually since all shRNA plasmids also contained a BFP marker.

### Plasmid Constructs

Fluorescent report constructs CaMKII synaptophysin-pHluorin, hDlxI56i synaptophysin-pHluorin, hDlxI56i iATPSnFR2.0-HALO (Bredvik and Ryan, 2024) and CaMKII iATPSnFR2.0-HALO (Marvin et al., 2024) are available on Addgene.

All shRNA plasmids were made by digesting pLKO.1 hPGK-BFP TRC cloning vector (Addgene #191566) with AgeI-HF and EcoRI-HF and annealed shRNA sequences were ligated with Quick Ligase Buffer (NEB). Primers for oligos against the listed sequences followed the format:

5’ CCGG-sense-CTCGAG-antisense-TTTTTG 3’, 5’AATTCAAAAA-sense-CTCGAG-antisense 3’

Target sequences:

BDH1: TTTGTGGGAAATACCTTTATA

MCU: GCTACCTTCTCGGCGAGAACGCTGCCAGTT

SCOT: GGAGGATCACCCATCAAATAT

### Virus production

HEK293FT cells were transfected using calcium phosphate transfection with lentiviral constructs and the associated packaging plasmids psPAX2 (a gift of Didier Trono, Addgene #12260) and pMD2.G (a gift of Didier Trono, Addgene 12259). 16 hours after transfection, culture media was exchanged for serum free viral production media: Ultraculture (Lonza), 1% (v/v) Penicillin-Streptomycin/L-glutamine, 1% (v/v) 100 mM sodium pyruvate, 1% (v/v) 7.5% sodium bicarbonate and 5 mM sodium butyrate. HEK293FT supernatants were collected at a timepoint of 46 hours after transfection and filtered through a 0.45 mm cellulose acetate filter. Viral supernatants were concentrated using Lenti-X Concentrator (Takara) and spun for 45 minutes at 1500 x g, then resuspended in PBS. These samples were aliquoted and stored at 80°C until use. Genomic titer was determined using a Lenti-X GoStix Plus kit (Takara). Lentivirus was functionally titrated previously using parallel preparation of GFP-expressing viral particles (Campeau et al., 2009) to determine the volume of lentivirus required to achieve ∼100% transduction of hippocampal and cortical neuron cultures, 106 GC (genome copies)/mL (Ashrafi et al., 2020). Lentivirus was added to neuron cultures at a timepoint of 2-4 days *in vitro*, and all experiments were performed at least 10 days after viral transduction to ensure adequate time for viral gene expression and protein knockdown.

### Statistical Analysis

GraphPad Prism v8 was used for statistical analysis and fitting, where the exponential decay function was used without constraints. For endocytic rates of fluorescence decay, data points starting 2 seconds after the end of stimulation were used in fitting. As specified in figure legends, the significance of differences between various conditions was calculated using nonparametric Mann-Whitney and Kruskal-Wallis tests for unpaired comparisons and Wilcoxon matched-pairs signed rank test for paired comparisons. We did not assume that the distributions of our datasets would follow a normal distribution. Generally, p < 0.05 was considered significant and denoted with one asterisk, while p < 0.01, p < 0.001, p < 0.0001 were denoted with two, three, or four asterisks, respectively. Bonferroni correction was used to correct for multiple comparisons in Kaplan-Meier analysis. Dunn’s correction was used to correct for multiple comparisons in all other analyses.

## Results

### Neuronal ketone metabolism supports presynaptic function

The metabolism of ketones by neurons is of interest as a potential therapy for various neurologic conditions with underlying metabolic perturbations, but it is unclear whether ketones themselves are metabolized in a cell autonomous fashion in neurons. Synaptic vesicle (SV) recycling rates are highly sensitive to metabolic perturbations and can be used as a surrogate for synaptic metabolism during and after electrical activity (Ashrafi et al., 2017; Ashrafi et al., 2020). Here, we made use of synaptophysin-pHluorin (physin-pHluorin) to target pHluorin to the lumen of SVs, using either a Ca^2+^/calmodulin-dependent protein kinase II (CaMKII) promoter to drive expression in excitatory neurons or a minimal beta globin promoter combined with the hDlxI56i enhancer (Mich et al., 2021; Dimidschstein et al., 2016) to drive expression in inhibitory neurons. We recently validated the specificity of this cell type specific expression scheme in our neonatal rat neuron primary culture system (Farrell et al., 2024).

To test the suitability of ketones as a presynaptic fuel, we examined the vesicle recycling response to a brief burst of action potentials (AP). During AP stimulation, the physin-pHluorin fluorescence signal rises abruptly, reflecting the balance of exocytosis and ongoing endocytosis, while following the stimulus period, the decay in fluorescence signal is dominated by a convolution of the endocytic recapture of physin-pHluorin and SV reacidification. We previously showed that acute block of ATP synthesis machinery leads to a rapid block of pHluorin recycling (Ashrafi et al., 2017; Ashrafi et al., 2020). Comparison of the SV endocytic time constant following a 100 AP stimulus in either 5 mM glucose or 5 mM β-hydroxybutyrate (BHB) in both excitatory and inhibitory nerve terminals showed that acute application of ketones provides a sufficient oxidizable carbon source to support SV recycling (Figure 1A, 1B, Supplementary Figure 1A-D). These experiments are carried out in co-cultures of astrocytes and neurons, leaving open the possibility that the critical ketone metabolism might be occurring in astrocytes that in turn under these conditions spare pyruvate produced through glycolysis and deliver it as lactate to neurons (Mächler et al., 2016; Roosterman and Cottrell, 2020). To determine whether neurons are self-reliant on ketone metabolism we decided to perturb ketone metabolism only in neurons. We designed shRNAs targeting two key mitochondrial enzymes required for ketone metabolism, β-hydroxybutyrate dehydrogenase 1 (BDH1) and succinyl-CoA:3-ketoacid CoA transferase (SCOT/OXCT1) (Figure 1C). The efficacy of knockdown (KD) was confirmed using Western blot of cortical neurons virally expressing the shRNA against BDH1 and SCOT (Figure 1D, 1E, Supplementary Figure 1E, 1F, Extended Data Figures 1-2, 1-4, 1-6, 1-8, S1-2, S1-4).

We used sparse transfection of both shRNA plasmids and physin-pHluorin in hippocampal neurons, selecting neurons expressing both plasmids for experiments (see Methods). In both excitatory and inhibitory neurons expressing the shRNA targeting BDH1, SV recycling kinetics were no different than controls when stimulated in glucose (Figure 1F, 1G, Supplementary Figure 1A-D). In contrast, when the same neurons were stimulated in the presence of BHB, endocytosis arrested immediately (Figure 1F, 1G). Similar results were obtained in cells expressing the shRNA targeting SCOT (Supplementary Figure 1G, 1H). In sum, these results demonstrate that the two neuronal subtypes studied here are both capable of catabolizing ketones and of using the ATP generated from ketolysis to fuel synaptic transmission, by confirming that BDH1 and SCOT are both necessary for neuronal ketone metabolism.

### Ketones maintain higher steady state levels of ATP in inhibitory compared to excitatory neurons

To directly compare the metabolic state of inhibitory and excitatory neurons using mitochondrial fuel sources, we targeted the recently improved ratiometric ATP sensor iATPSnFR2.0-HALO (Marvin et al., 2024) to the cytosol of either excitatory or inhibitory neurons using CaMKII and hDlxI56i specific expression constructs. We recently showed that, in agreement with *in vivo* findings (Nie and Wong-Riley, 1995; Gulyás et al., 2006), cultured inhibitory axons have a ∼25% higher density of mitochondria compared to excitatory axons, resulting in much higher resting ATP levels when axons are fueled solely by mitochondrial metabolism using a mixture of lactate and pyruvate (Bredvik and Ryan, 2024). Following 5 minutes of perfusion in BHB, we found that resting ATP levels were ∼3-fold higher in inhibitory compared to excitatory axons (Figure 2A), consistent with the notion that neuronal mitochondria carry out ketolysis (Figure 1) and that inhibitory neurons have a greater total mitochondrial capacity than excitatory neurons.

**Figure 2.**
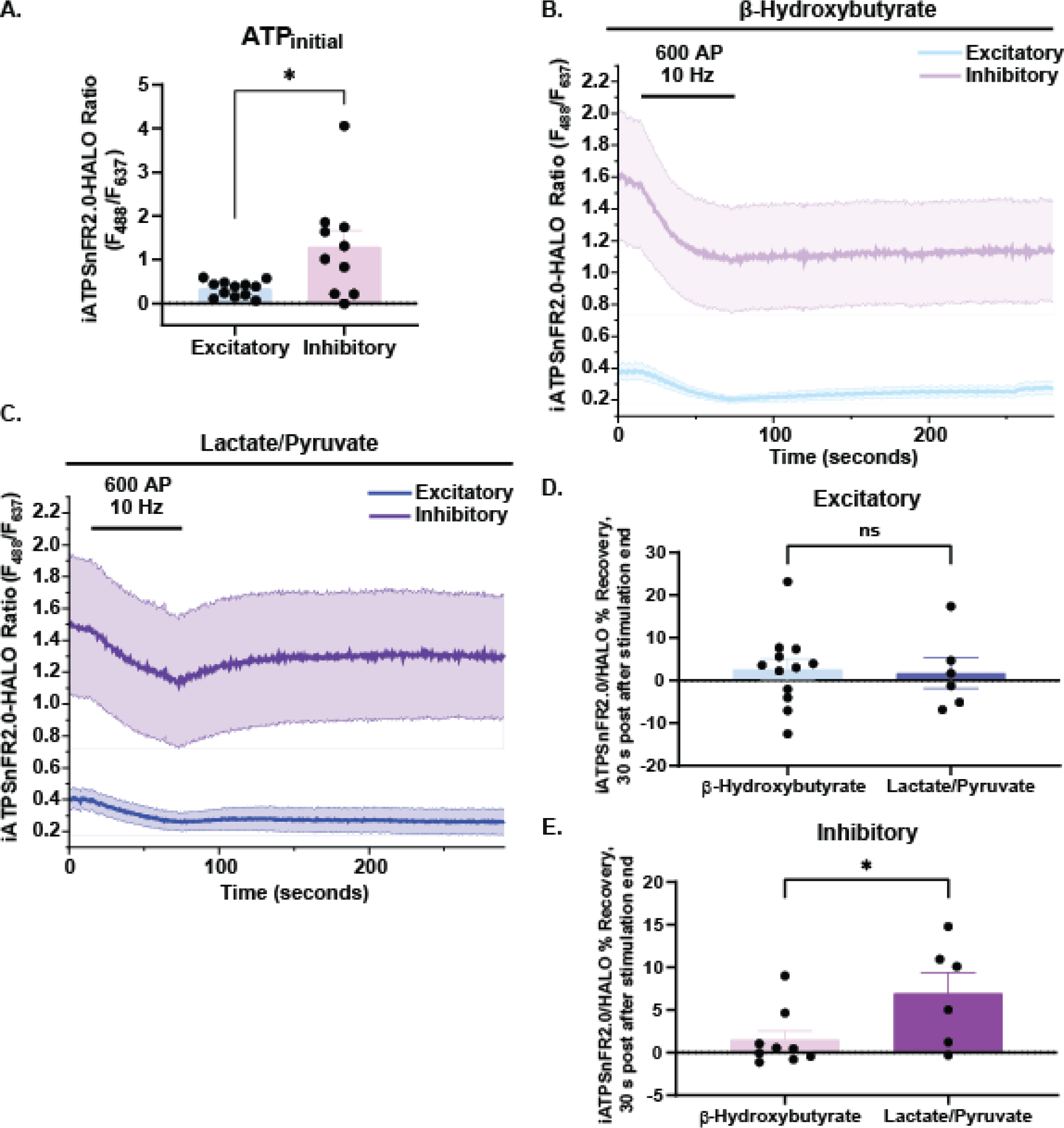
Inhibitory neurons using lactate/pyruvate as fuel perform better than those relying on β-hydroxybutyrate. **A)** Quantification of iATPSnFR2.0/HALO ratio in excitatory (CaMKII) and inhibitory (hDlxI56i) neurons prior to stimulation in β-hydroxybutyrate (n=12 excitatory, mean=0.3371; n=10 inhibitory, mean=1.288; Error bars SEM, *p=0.0358, Mann-Whitney test). **B)** Fluorescence ratio traces of excitatory and inhibitory neurons in β-hydroxybutyrate during stimulation with 600 action potentials (AP) at 10 Hz (n=12 excitatory, n=10 inhibitory, error bands SEM). **C)** Fluorescence ratio traces of excitatory and inhibitory neurons in lactate/pyruvate during stimulation with 600 AP at 10 Hz (n=5 excitatory, n=5 inhibitory, error bands SEM). **D)** Fluorescence ratio recovery in excitatory neurons following stimulation in lactate/pyruvate or β-hydroxybutyrate (n=6 lactate/pyruvate, mean=1.736 %; n=12 β-hydroxybutyrate, mean=2.554 %; Error bars SEM, n.s., p=0.682, Mann-Whitney test). **E)** Fluorescence ratio recovery in inhibitory neurons following stimulation (n=6 lactate/pyruvate, mean=6.949 %; n=9 β-hydroxybutyrate, mean=1.463 %; Error bars SEM, *p=0.036, Mann-Whitney test).

Given the similarity of resting ATP levels between neurons in BHB and neurons in lactate/pyruvate (Figure 2A, Supplementary Figure 2A), we examined iATPSnFR2.0-HALO fluorescence recovery following a burst of AP firing capable of transiently depressing presynaptic ATP levels, for neurons bathed in either BHB or lactate/pyruvate. Following a 60 second burst of 10 Hz AP firing, presynaptic ATP recovered much more slowly in excitatory compared to inhibitory neurons when fueled by lactate/pyruvate (Figure 2C-E), as previously reported (Bredvik and Ryan, 2024). Inhibitory neurons in BHB recovered much more slowly than inhibitory neurons in lactate/pyruvate (Figure 2B-2D). Remarkably, when fueled by BHB, the kinetics of ATP recovery are much slower in both neuron types (Figure 2B, 2D-E). These data suggest that although steady state bioenergetic metabolism of ketones results in significantly higher resting presynaptic ATP levels in inhibitory neurons, the details of regulation for ketolysis during activity appear to be weaker or less efficient than for oxidative combustion of lactate and pyruvate, resulting in a longer delay in regenerating ATP under conditions of ketone metabolism.

### Excitatory and inhibitory neurons robustly support synaptic transmission with different mitochondrial fuels

To assess how differences in baseline ATP levels and recovery affected synaptic transmission in excitatory and inhibitory neurons, physin-pHluorin was expressed in both neuron types. Since the endocytosis/reacidification step of the SV cycle is very sensitive to metabolic perturbations (Ashrafi et al., 2017, Ashrafi et al., 2020), we characterized the kinetics of this step during endocytic recovery of SVs after stimulation using this pHluorin-tagged SV protein. We used two different stimulus paradigms to examine whether synapses running on ketolysis would show any impact on SV recycling. First, using sustained 10 Hz firing to transiently deplete ATP (Figure 3B, 3C) we examined the physin-pHluorin fluorescence recovery rate after stimulation by normalizing average traces to peak fluorescence and plotting them on a semi-log plot (Supplementary Figure 2B, 2C). Neurons in either lactate/pyruvate or BHB were able to recover fluorescence after stimulation, indicating that the bioenergetic state of neurons using mitochondrial fuel alone was sufficient to fuel the processes of endocytosis and reacidification in an acute setting. We did not observe any differences in time constants of endocytosis or maximal fluorescence between excitatory and inhibitory neurons in either fuel (Supplementary Figure 2D, Supplementary Table 1).

**Figure 3.**
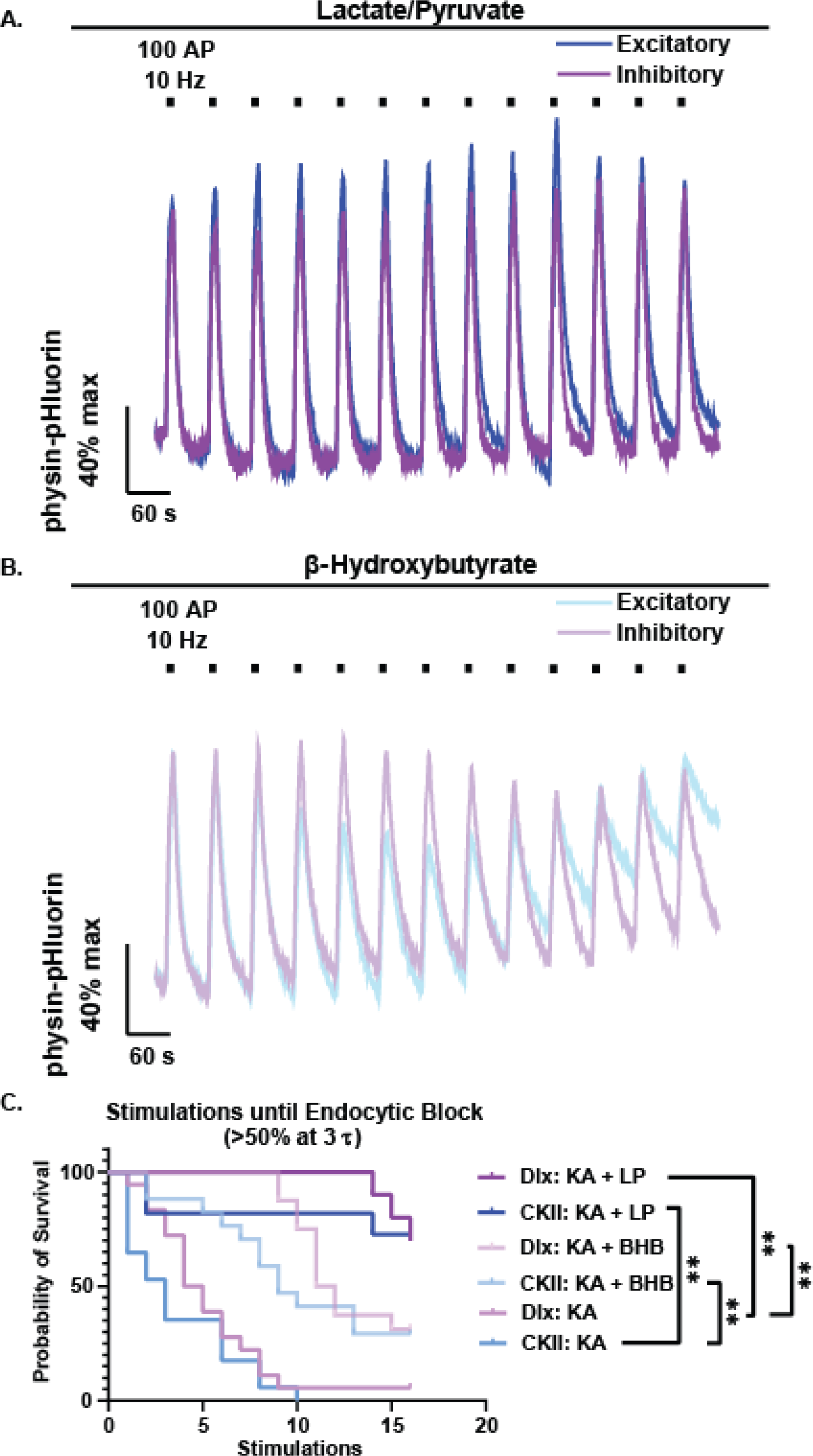
Excitatory and inhibitory neurons perform vesicle recycling better in lactate/pyruvate than in β-hydroxybutyrate. **A)** Representative traces of excitatory and inhibitory neurons expressing synaptophysin-pHluorin (physin-pHluorin) stimulated with 100 action potentials (AP) every minute at 10 Hz, fueled by lactate and pyruvate (LP; 1.25 mM each) in presence of glycolytic inhibitor koningic acid (KA). **B)** Representative traces of excitatory and inhibitory neurons fueled by β-hydroxybutyrate (BHB; 5 mM racemic mixture) with 100 AP, 10 Hz stimulation in KA and glucose. **C)** Survival analysis of number of 100 AP, 10 Hz stimulations before reaching endocytic block of > 50% at the specified timepoint of 3 endocytic time constants after stimulation (n=18 inhibitory KA, median survival=4.5 stimulations; n=16 excitatory KA, median survival=3 stimulations; n=16 inhibitory BHB/gluc+KA, median survival=11.5 stimulations; n=17 excitatory BHB/gluc+KA, median survival=9 stimulations; n=10 inhibitory LP/gluc+KA, median survival=undefined/16+ stimulations; n=11 excitatory LP/gluc+KA, median survival=undefined/16+ stimulations; **p<0.005, Kaplan-Meier analysis).

In a second approach, we used a synaptic endurance assay that we recently developed where neurons are subjected to repeated rounds of stimulation at minute intervals (Kokotos et al., 2023). We previously showed that under sufficient glycolytic fuel conditions, excitatory neurons are capable of sustaining at least ten rounds of stimulation without a significant slowing of SV recycling (Kokotos et al., 2023). Here, we found that both excitatory and inhibitory neurons can sustain synaptic function under this paradigm when fueled by either lactate and pyruvate or BHB (Figure 3A, 3B). As a metric for characterizing the ability of different neuron types to sustain synapse function under repeated stimulation, we made use of a survival analysis on a cell-by-cell basis where we defined survival as the number of stimulations where SV recycling resulted in at least 50% fluorescence recovery within 3 time constants of the control endocytic recovery (defined by the same cell’s recovery in glucose; Pulido and Ryan, 2021). As a positive control, we show that when provided only glucose as fuel, both neuron types arrest synapse function on average by approximately the 7^th^ round of stimulation in the presence of the GAPDH inhibitor koningic acid (KA). Consistent with the faster ATP recovery kinetics in lactate/pyruvate compared to BHB, we found that both neuron types survived longer in the synapse endurance paradigm in lactate/pyruvate compared to BHB (Figure 3C). Notably, this approach did not reveal any significant differences in the ability to sustain synapse function in excitatory versus inhibitory neurons in either of the mitochondrial fuels tested (Supplementary Figure 3).

### Efficient ketolysis in presynaptic mitochondria requires mitochondrial Ca^2+^ uptake

We previously showed that axonal mitochondrial metabolism depends on mitochondrial Ca^2+^ uptake via the mitochondrial Ca^2+^ uniporter (MCU) to upregulate presynaptic mitochondria-based ATP production during electrical activity (Ashrafi et al., 2020). This Ca^2+^-dependent metabolic regulation is thought to be mediated by upregulation of the activity of PDH and the TCA cycle dehydrogenases isocitrate dehydrogenase and oxoglutarate dehydrogenase (Denton, 2009). While pyruvate metabolism would in principle rely on activation of all three of these enzymes, ketolysis would rely only on the TCA cycle dehydrogenases (Figure 4A). At present the relative importance of these different steps of Ca^2+^ control of mitochondrial metabolism is unknown, but one possible reason for the kinetics of ATP production following stimulation in BHB compared to pyruvate being less efficient is that regulation of PDH activity by Ca^2+^ predominates in the overall upregulation of mitochondrial ATP production. To test this idea, we reasoned that synaptic function under BHB metabolism might be insensitive to loss of MCU, and therefore examined synapse function using physin-pHluorin in both neuron types following MCU KD in glucose, lactate/pyruvate or BHB. Loss of MCU in both excitatory and inhibitory neurons led to a rapid arrest of SV recycling when neurons were fueled with lactate/pyruvate, while MCU KD had no impact on synapse function when neurons relied on glucose as the primary fuel (Figure 4B, 4D), consistent with our previous findings in excitatory neurons (Ashrafi et al., 2020). Remarkably, both neuron types were also highly sensitive to loss of MCU when fueled purely by BHB (Figure 4C, 4E), implying that other metabolic steps separate from PDH rely on efficient mitochondrial Ca^2+^ uptake.

**Figure 4.**
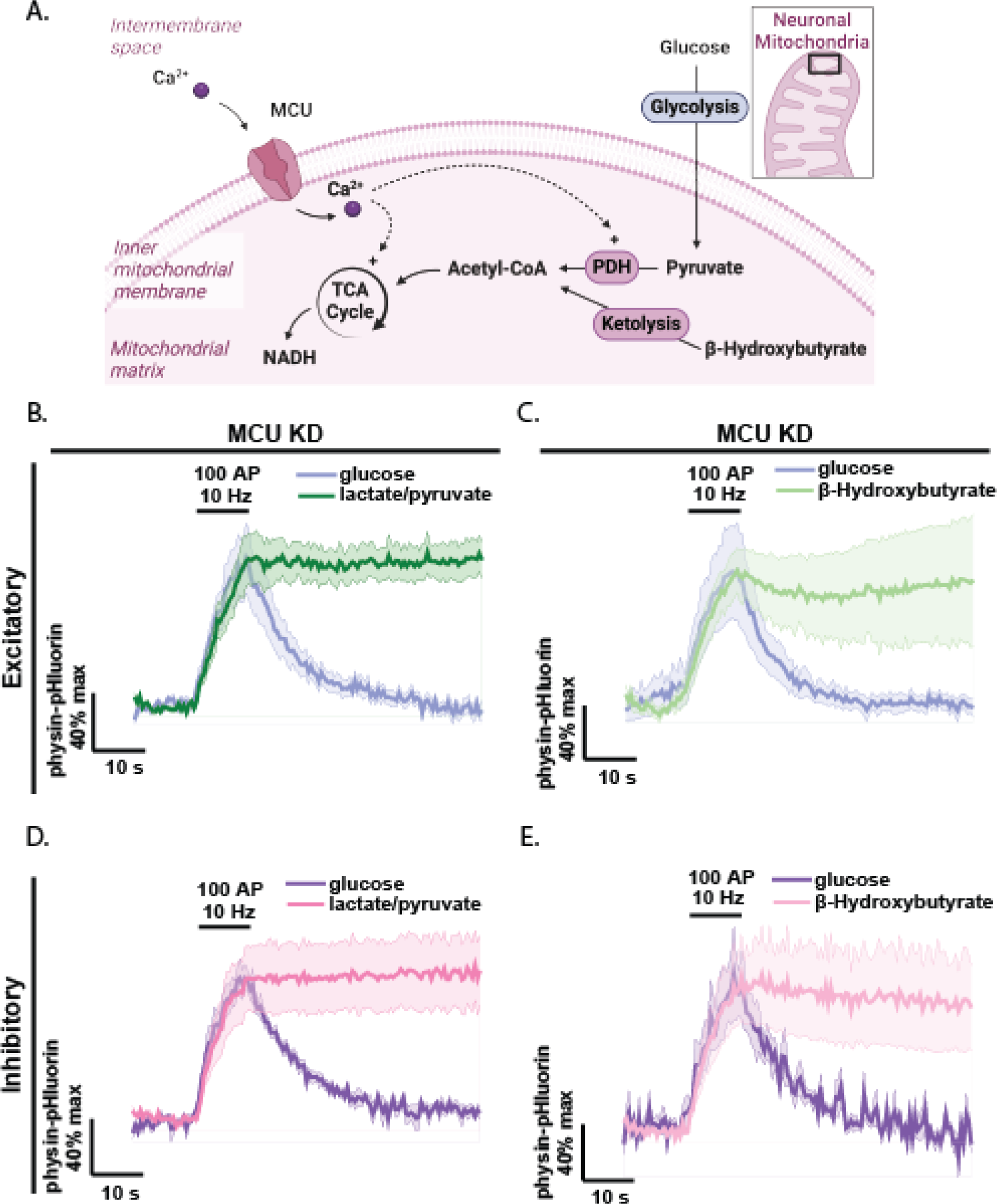
Pyruvate dehydrogenase activation by Ca^2+^ is a key point of mitochondrial metabolic regulation in active excitatory and inhibitory neurons. **A)** Diagram of activity dependent upregulation of mitochondrial metabolism. **B-C)** Fluorescence traces of excitatory neurons expressing synaptophysin-pHluorin (physin-pHluorin) and MCU shRNA stimulated with 100 action potentials (AP) at 10 Hz, in presence of 5 mM glucose followed by 10 µM koningic acid (KA) and either **B)** 1.25 mM lactate and 1.25 mM pyruvate (n=7, error bands SEM. Data pooled with neurons in lactate/pyruvate without glucose and KA) or **C)** 5 mM β-hydroxybutyrate (n=7, error bands SEM). **D-E)** Fluorescence traces of inhibitory neurons expressing physin-pHluorin and MCU shRNA, stimulated with 100 AP at 10 Hz in 5 mM glucose followed by 10 µM KA and either **D)** 1.25 mM lactate and 1.25 mM pyruvate (n=6 MCU KD, error bands SEM. Data pooled with neurons in lactate/pyruvate without glucose and KA) or **E)** 5 mM β-hydroxybutyrate (n=8 MCU KD, error bands SEM).

## Discussion

Here, we show that hippocampal inhibitory and excitatory neurons are capable of metabolizing ketones at sufficient rates to fuel moderate amounts of synaptic activity, and that this metabolism is mediated cell autonomously through the ketolytic enzymes BDH1 and SCOT. With the newly improved ATP sensor iATPSnFNR2.0-HALO, we show that active inhibitory neurons produce ATP from pyruvate more rapidly than from BHB, while excitatory neurons do not. Lastly, we use comparisons between ketone and pyruvate metabolism to identify largely similar Ca^2+^-dependent mechanisms of upregulating mitochondrial metabolism in excitatory and inhibitory neurons through the TCA cycle.

Ketones are canonically considered a fuel for the fasting brain when glucose is limited, but to our knowledge it has not yet been shown that metabolism of ketones can occur efficiently, cell autonomously, and in specific subtypes of neurons. Excitatory and inhibitory subtypes of neurons differentially express genes related to mitochondrial metabolism (Wynne et al., 2021) and have different functional ATP requirements due to their different intrinsic firing rates (Kann et al., 2011). It is therefore possible that the metabolism of ketones could vary between neuronal subtypes in physiologically significant ways. Here, we sought to conclusively determine whether mammalian excitatory and inhibitory neurons could use the ketone BHB as fuel for synaptic activity, and whether inhibitory and excitatory neurons performed ketone metabolism with differing efficiency.

One notable finding is that under pure BHB conditions, inhibitory neurons sustain much higher steady-state presynaptic ATP levels than excitatory neurons (Figure 2A), consistent with our previous observation that there is a higher abundance of mitochondria in inhibitory neurons (Bredvik and Ryan, 2024). We found that although both subtypes could cell autonomously use BHB as a fuel for synaptic transmission, in the stimulus paradigms examined (10 Hz firing in bursts at minute intervals or sustained for 60 seconds) inhibitory neurons performed no better than excitatory neurons when provided BHB as fuel (Figure 1, Figure 3, Supplementary Figure 1, Supplementary Figure 2). Since ATP replenishment following activity is slower for excitatory neurons than for inhibitory neurons in lactate and pyruvate conditions and both neuron types have slower ATP recovery in BHB following activity, the specific stimulus paradigms we examined may not be sufficiently taxing for synaptic transmission to be sensitive to these differences in ATP metabolism. It is also possible that *in vivo* changes in other brain metabolites in combination with ketones might drive differences in changes in synaptic function across these neuron types. For example, brain lactate has been observed to increase with fasting in tandem with BHB (Pan et al., 2000), indicating that the body is able to provide these fuels to neurons in conditions of lower glucose availability. It remains possible that the metabolic perturbations of the ketogenic diet have minimal effects on synaptic transmission, given the many other proposed mechanisms of ketone action (Masino et al., 2011; Youm et al., 2015; Qiao et al., 2024; Ma et al., 2007; Juge et al., 2010), but our recent results highlight a previously unconsidered therapeutic mechanism. Whether brain lactate levels similarly increase in prolonged ketogenic diet use represents an interesting topic for future study.

Since using BHB as fuel could be interpreted as lessening the apparent advantage of inhibitory neuronal metabolism over that of excitatory neurons in our ATP measurements (Figure 2), PDH could be a key differentially regulated step in inhibitory neuron oxidative metabolism. This hypothesis is consistent with a recent publication that finds a positive correlation between phosphorylation of PDH and neuronal firing rate (Yang et al., 2024), since inhibitory neurons tend to have higher firing rates. The lower efficiency of BHB as a fuel for nerve terminals could alternatively be due to insufficient baseline expression of ketolytic enzymes for rapid ATP production during synaptic transmission. While BHB was able to support synaptic transmission in the short term, it was less efficient than an equimolar amount of lactate and pyruvate and there was no difference in the performance of excitatory and inhibitory neurons using BHB as fuel (Figure 3). Since the metabolic benefit of ketones is predicated on better endurance of inhibitory neurotransmission in comparison to excitatory neurotransmission when BHB is used as a fuel source, we did not expect to find that neither of the stimulus paradigms we used could reveal a difference in synaptic endurance between neuron types. Although our results are consistent with differential mitochondrial metabolism of pyruvate in inhibitory neurons with limited glucose, ketones specifically seemed to fuel excitatory and inhibitory synaptic activity similarly in an acute setting.

While we performed shRNA-mediated knockdown rather than genetic knockout of ketolytic enzymes, endocytic arrest remarkably occurred immediately in nerve terminals stimulated in the presence of BHB (Figure 1; Supplementary Figure 1). In contrast, nerve terminals in BHB had sufficient ATP to be able to sustain heavy stimulation and the concomitant endocytosis and vesicle acidification required for the observed drop in fluorescence (Supplementary Figure 2). Even in the presence of glycolytic block (KA) with only glucose provided as fuel, neurons were able to perform multiple rounds of stimulation (Figure 3C). These results suggest that neurons with glycolytic block have the capacity to shift to using alternative sources of fuel for short periods of time, such as stored glycolytic intermediates, ketones or lactate provided by neighboring astrocytes (Mächler et al., 2016; Thevenet et al., 2016). Since eliminating ketone metabolism seems to eliminate much of this ability, we posit that ketones could be a primary source of this metabolic flexibility. Astrocytes may pass BHB to nearby neurons, particularly in the condition of a high fat diet (Thevenet et al., 2016; Guzmán and Blázquez, 2001; Silva et al., 2022), and this hypothesized pathway could be investigated by providing glucose and KA to BDH1 knockdown neurons in future experiments.

In this study, we provided a relatively high effective concentration of BHB that approximates the blood concentration of ketones in patients on a ketogenic diet (van Delft et al., 2010), but overestimates the concentration of ketones encountered by brain tissue (Pan et al., 2000). Our experiments therefore should be sensitive to any effects of differential enzyme abundance between excitatory and inhibitory neurons, since BHB is constantly perfused directly onto neurons in excess. However, since these experiments occurred on an acute time scale of minutes there was presumably insufficient time for gene expression to change in response to the changing metabolic conditions. Longer time courses as well as studies of differential gene expression and metabolomic changes in neurons metabolizing ketones will more conclusively determine the role for metabolism in the mechanism of the ketogenic diet in treating epilepsy.

## Author Contributions

K.B. and T.A.R. contributed to the experimental design. K.B. and C.L. collected and analyzed the data. K.B. and T.A.R. contributed to writing the manuscript.

## Declaration of Interests

The authors have no conflicts of interest to disclose.

## Acknowledgements

This work was supported by NIH grant 2R01NS036942 awarded to T.A.R. and by a Medical Scientist Training Program grant from the National Institute of General Medical Sciences of the NIH under award number T32GM007739 to the Weill Cornell/Rockefeller/Sloan Kettering Tri-Institutional MD-PhD Program. Some figures were made with Biorender. We thank Giulia Dalaty and Hayoung Lee for technical assistance, and we thank members of the Ryan lab for helpful discussion, especially Ryan J. Farrell, Ghazaleh Ashrafi and Jaime de Juan-Sanz for essential advice.

## Notes

### Competing Interest Statement

The authors have declared no competing interest.

