## Supplemental Figures for "Characterization of β-Hydroxybutyrate as a Cell Autonomous Fuel for Active Excitatory and Inhibitory Neurons"

**Supplementary Figure 1.**

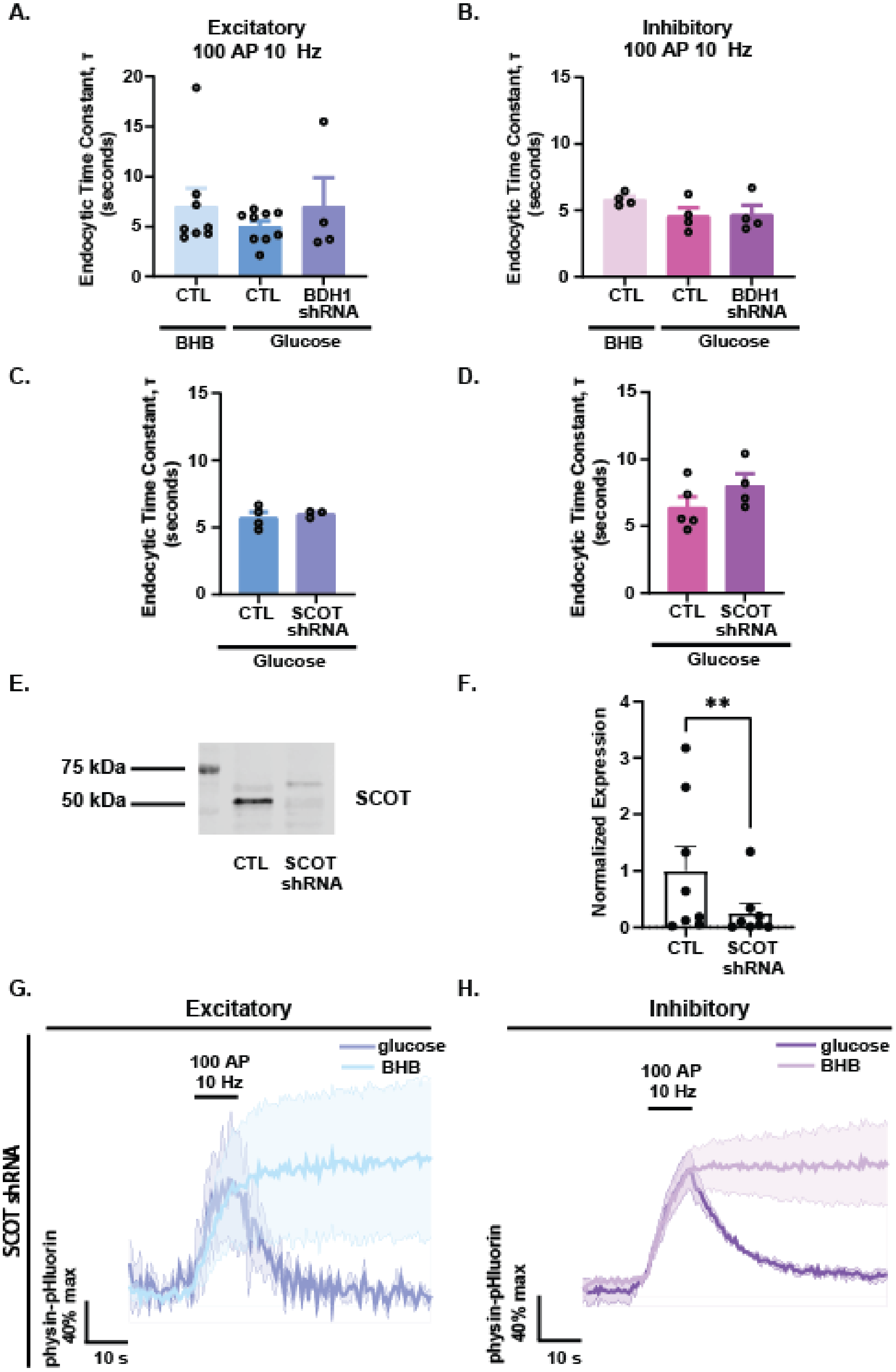

**Supplementary Figure 2.**

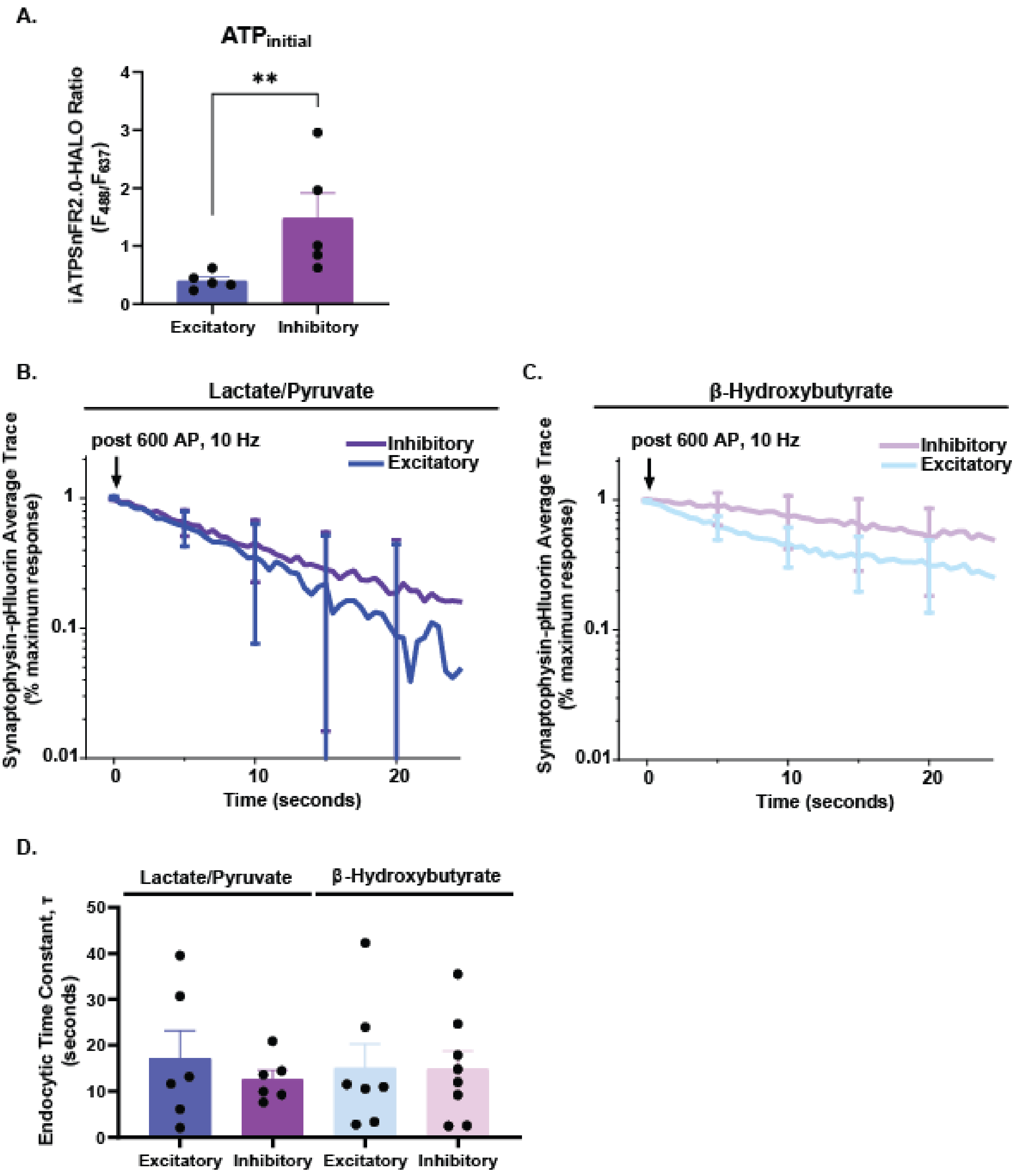

**Supplementary Figure 3.**

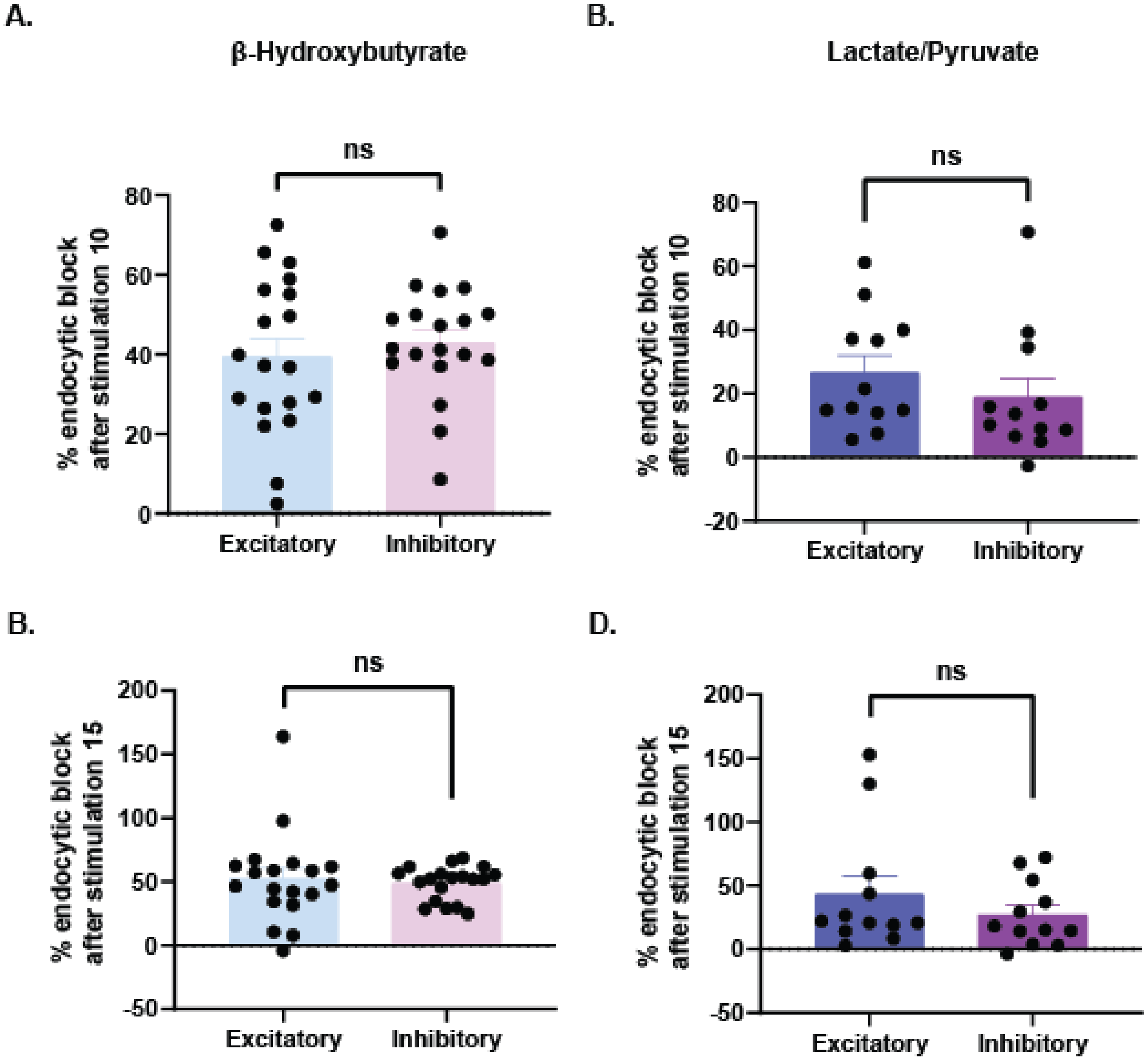

### Supplementary Table 1.

**Excitatory**

| BDH1 shRNA gluc | BDH1 shRNA BHB | SCOT shRNA gluc | SCOT shRNA BHB | LP gluc KA | BHB gluc KA | gluc KA | CTL LP | MCU shRNA LP | CTL BHB | MCU shRNA BHB | BHB gluc KA 600 AP | LP gluc KA 600 AP |
|---|---|---|---|---|---|---|---|---|---|---|---|---|
| 0.1269 | 0.1055 | 0.06403 | 0.1256 | 0.1469 | 0.07606 | 0.1318 | 0.1598 | 0.1326 | 0.05012 | 0.02360 | 0.09563 | 0.2765 |
| 0.05075 | 0.00127 | 0.01269 | 0.03583 | 0.2312 | 0.02283 | 0.1010 | 0.1063 | 0.07401 | 0.12244 | 0.05582 | 0.02512 | 0.01480 |
| 0.07853 | 0.2702 | 0.0127 | 0.01888 | 0.02825 | 0.07857 | 0.1111 | 0.07884 | 0.1588 | 0.06259 | 0.04528 | 0.1306 | 0.03351 |
| 0.1149 | 0.02233 | | | 0.06481 | 0.1294 | 0.1042 | 0.1313 | 0.1809 | 0.04032 | 0.08386 | 0.01615 | 0.02979 |
| | | | | 0.2325 | 0.1682 | 0.08102 | 0.05404 | 0.2275 | 0.06776 | 0.02010 | 0.02277 | 0.09576 |
| | | | | 0.08284 | 0.1800 | 0.06484 | 0.1039 | 0.06533 | 0.05000 | 0.0366 | 0.02729 | 0.02681 |
| | | | | 0.07567 | 0.1135 | 0.12811 | | 0.09237 | | 0.07571 | 0.08647 | |
| | | | | 0.1472 | 0.06732 | 0.04473 | | | | | | |
| | | | | 0.06495 | 0.09420 | 0.1042 | | | | | | |
| | | | | 0.1074 | 0.10538 | 0.04878 | | | | | | |
| | | | | 0.03677 | 0.1424 | 0.2081 | | | | | | |
| | | | | | 0.02786 | 0.10173 | | | | | | |
| | | | | | 0.03249 | 0.06981 | | | | | | |
| | | | | | 0.01210 | 0.1344 | | | | | | |
| | | | | | 0.08852 | 0.07428 | | | | | | |
| | | | | | 0.1940 | 0.04359 | | | | | | |
| | | | | | 0.1743 | | | | | | | |
| | | | | | 0.05279 | | | | | | | |

**Inhibitory**

| BDH1 shRNA gluc | BDH1 shRNA BHB | SCOT shRNA gluc | SCOT shRNA BHB | LP gluc KA | BHB gluc KA | gluc KA | CTL LP | MCU shRNA LP | CTL BHB | MCU shRNA BHB | BHB gluc KA 600 AP | LP gluc KA 600 AP |
|---|---|---|---|---|---|---|---|---|---|---|---|---|
| 0.04605 | 0.1288 | 0.07466 | 0.1147 | 0.05672 | 0.07283 | 0.08191 | 0.06958 | 0.0725 | 0.04614 | 0.0681 | 0.01371 | 0.06731 |
| 0.04129 | 0.01386 | 0.1309 | 0.09364 | 0.1199 | 0.04442 | 0.1449 | 0.02326 | 6.667E-05 | 0.03964 | 0.3100 | 0.04474 | 0.01565 |
| 0.1081 | 0.2552 | 0.07249 | 0.08340 | 0.04688 | 0.1590 | 0.1721 | 0.08118 | 0.4243 | 0.13429 | 0.0681 | 0.05374 | 0.02522 |
| 0.07939 | 0.3749 | 0.06570 | 0.04104 | 0.06695 | 0.1539 | 0.2193 | 0.07315 | 0.2655 | 0.0849 | 0.1391 | 0.01566 | 0.02868 |
| | | | | 0.1510 | 0.1550 | 0.3436 | | 0.1130 | | 0.08750 | 0.01956 | 0.09015 |
| | | | | 0.09359 | 0.0608 | 0.1378 | | 0.2527 | | 0.1179 | 0.03164 | 0.01446 |
| | | | | 0.1017 | 0.1480 | 0.1427 | | 0.2030 | | 0.04342 | 0.05129 | |
| | | | | 0.09394 | 0.08212 | 0.14043 | | | | 0.05003 | 0.009106 | |
| | | | | 0.1520 | 0.06281 | 0.09519 | | | | | 0.04177 | |
| | | | | 0.08745 | 0.2751 | 0.05807 | | | | | | |
| | | | | | 0.1895 | 0.09833 | | | | | | |
| | | | | | 0.03789 | 0.1109 | | | | | | |
| | | | | | 0.03862 | 0.01571 | | | | | | |
| | | | | | 0.01321 | 0.08698 | | | | | | |
| | | | | | 0.1014 | 0.1153 | | | | | | |
| | | | | | 0.08448 | 0.1813 | | | | | | |
| | | | | | 0.1472 | | | | | | | |

### Supplementary Figure Legends

**Supplementary Figure 1. Perturbations of ketone metabolism affect synaptic vesicle recycling in neurons fueled with BHB, but not those fueled with glucose. A)** Quantification of time constants from excitatory neurons transfected with synaptophysin-pHluorin (physin-pHluorin) and shRNA against β-hydroxybutyrate dehydrogenase 1 (BDH1) with 100 action potential (AP), 10 Hz stimulation in glucose (n=8 CTL BHB, mean=7.065 s; n=9 CTL glucose, mean=5.036 s; n=4 BDH1 sh, mean=7.032 s; error bars SEM, all comparisons n.s., Kruskal-Wallis test). **B)** Time constants from inhibitory neurons transfected with physin-pHluorin and BDH1 shRNA with 100 AP, 10 Hz stimulation in glucose (n=4 CTL BHB, mean=5.836 s; n=4 CTL glucose, mean=4.6 s; n=4 BDH1 sh, mean=4.692 s. Error bars SEM, all comparisons n.s., Kruskal-Wallis test). **C)** Time constants from excitatory neurons transfected with physin-pHluorin and succinyl-CoA:3-ketoacid CoA transferase (SCOT) shRNA with 100 AP, 10 Hz stimulation in glucose (n=4 CTL, mean=5.737 s; 3 SCOT sh, mean=6.008 s; error bars SEM). **D)** Time constants from inhibitory neurons transfected with physin-pHluorin and SCOT shRNA with 100 AP, 10 Hz stimulation in glucose (n=5 CTL, mean=6.424 s; 4 SCOT sh, mean=8.041 s; error bars SEM). **E)** Representative Western blot against succinyl-CoA:3-ketoacid CoA transferase (SCOT) WT cortical cultures and cultures treated with SCOT shRNA virus for 10 days (full blots in Extended Data Figure S1-1, S1-2, S1-3, S1-4). **F)** Quantification of SCOT knockdown efficiency in Western blot, normalized to average signal of uninfected controls (N=4 cultures, n=2 technical replicates, mean=0.2533. Error bars SEM, **p=0.0078, Wilcoxon matched-pairs signed rank test). **G)** Synaptic vesicle cycling traces in excitatory neurons expressing CaMKII synaptophysin-pHluorin (physin-pHluorin) and shRNA against SCOT, in glucose or β-hydroxybutyrate (BHB) with glucose and glycolytic inhibitor koningic acid (KA) during 100 action potential (AP), 10 Hz stimulus (n=3, error bands SEM). **H)** Same as G), but with inhibitory neurons expressing hDlxI56i physin-pHluorin (n=4, error bands SEM).

**Supplementary Figure 2. Excitatory and inhibitory neurons recycle synaptic vesicles while using mitochondrial fuel sources. A)** Quantification of iATPSnFR2.0/HALO ratio prior to stimulation in lactate/pyruvate (n=5 excitatory, mean=0.4011; n=5 inhibitory, mean=1.481. Error bars SEM, **p=0.0079, Mann-Whitney test). **B)** Traces of inhibitory neurons expressing hDlxI56i synaptophysin-pHluorin (physin-pHluorin) and excitatory neurons expressing CaMKII physin-pHluorin stimulated with 600 action potentials (AP) at 10 Hz while in the presence of lactate/pyruvate (LP; 1.25 mM each), koningic acid (KA) and 5 mM glucose (LP+KA; n=6 excitatory, n=6 inhibitory, error bars SEM). **C)** Traces of inhibitory neurons and excitatory neurons expressing physin-pHluorin stimulated with 600 AP at 10 Hz while in the presence of β-hydroxybutyrate (BHB; 5 mM racemic mixture), koningic acid and 5 mM glucose (BHB+KA; n=8 excitatory, n=8 inhibitory, error bars SEM). **D)** Time constants of synaptic vesicle endocytosis in excitatory and inhibitory neurons using LP or BHB as fuel, from traces in A) and B) (n=6 excitatory LP+KA, mean=17.20 s; n=6 inhibitory LP+KA, mean=12.62 s; n=7 excitatory BHB+KA, mean=15.06 s; n=8 inhibitory BHB+KA, mean=14.87 s. Error bars SEM, all comparisons n.s., Kruskal-Wallis test).

**Supplementary Figure 3. Excitatory and inhibitory neurons recycle synaptic vesicles to similar extent after repeated stimulation in ketones. A)** Quantification of percent endocytic block after tenth 100 action potential (AP), 10 Hz stimulation in excitatory and inhibitory neurons expressing synaptophysin-pHluorin (physin-pHluorin) and fueled by β-hydroxybutyrate (n=19 excitatory, mean=39.58%; n=19 inhibitory, mean=43.1%. Error bars SEM. n.s., p=0.4879, Mann-Whitney test). **B**) Quantification of percent endocytic block after tenth 100 AP, 10 Hz stimulation in neurons expressing physin-pHluorin and fueled by lactate and pyruvate (n=12 excitatory, mean=26.62%; n=12 inhibitory, mean=18.95%. Error bars SEM. n.s., p=0.1978,

Mann-Whitney test). **C)** Quantification of percent endocytic block after fifteenth 100 AP, 10 Hz stimulation in neurons fueled by β-hydroxybutyrate (n=19 excitatory, mean=52.18%; n=19 inhibitory, mean=48.99%. Error bars SEM. n.s., p=0.9081, Mann-Whitney test). **D)** Quantification of percent endocytic block after fifteenth 100 AP, 10 Hz stimulation in neurons fueled by lactate and pyruvate (n=12 excitatory, mean=43.4%; n=12 inhibitory, mean=27.2%. Error bars SEM. n.s., p=0.4428, Mann-Whitney test).

**Supplementary Table 1. Maximum fluorescence of synaptophysin-pHluorin traces.** Fluorescence values from normalized excitatory (above) and inhibitory (below) neuron synaptophysin-pHluorin traces in main and supplemental figures reported here as absolute change in fluorescence with stimulation as a fraction of maximum possible fluorescence in presence of ammonium chloride at pH 7.4. All values truncated to 4 significant figures.
